# Body size cascades from *Neodiprion* sawfly hosts to *Perilampid* parasitoids: An investigation of sequential divergence

**DOI:** 10.64898/2026.11.04.764306

**Authors:** Sarah V. White, Salahuddin Mohammed, Robin K. Bagley

## Abstract

Ecological speciation drives diversity in plant-eating insect systems, but it may also generate parallel specialization in parasitoid associates in a cascading effect of “sequential divergence.” However, sequential divergence has never been successfully identified in parasitoids of externally feeding insects. To address this gap, we consider the *Pinus-Neodiprion-Perilampus neodiprioni* system, where plant-associated, externally feeding *Neodiprion* demonstrate ecological divergence, as supported by body size as a magic trait in sympatric species. It seems possible that this body size differentiation may cascade to *Neodiprion’s* parasitoids via sequential divergence. Therefore, we asked, “Does *P. neodiprioni* exhibit host-specific divergence in body size that parallels patterns in *Neodiprion* hosts?” Further, “Does *P. neodiprioni* exhibit host-related variation in the abdominal region or sexually dimorphic features, supporting reproductive barriers?” Using by-catch samples, we measured body size of *P. neodiprioni,* reared from *Neodiprion lecontei* and *Neodiprion pinetum* hosts. We analyzed body size variation as predicted by host species, parasitoid sex, host sex, host cocoon size, latitude, and their interactions. In addition, we used mixed models to analyze the effects of parasitoid sex and *Neodiprion* host on three size-based facial features. We found that *P. neodiprioni* demonstrates host-associated body size variation that mirrors size differences in their *Neodiprion* hosts. In addition, host-associated size variation occurs in the abdominal region and two sexually dimorphic characters of *P. neodiprioni*, supporting possible reproductive isolation. Together, these patterns suggest that host magic traits may drive not only within-species ecological divergence, but they can facilitate parasitoids’ subsequent morphological differentiation and possible sequential divergence.

## Introduction

During ecological speciation, populations in different environments experience divergent natural selection, which incidentally creates barriers to gene flow (Nosil, 2012; Rundle & Nosil, 2005). Thus, adaptive divergence can lead to reproductive isolation between populations, facilitating speciation (Elias et al., 2012). This phenomenon has been well-documented across many taxa, including South Atlantic island finches (Ryan et al., 2007), postglacial fishes (Schluter, 1996), and blind mole rats (Jiao et al., 2021), but it is perhaps most studied in plant-eating insect systems (Elias et al., 2012; Jean & Jean-Christophe, 2010; Nosil, 2004).

In plant-herbivore systems, distinct host plants serve as different environments, so host shifts can lead to ecological speciation (Feder & Forbes, 2010). Different traits may be favored on each host plant, reducing opportunities for gene flow between insect populations across plants. In particular, many insect systems display extreme habitat preference and host fidelity, mating on the host plants where they feed. This results in habitat isolation, a form of reproductive isolation (Feder & Forbes, 2010; Jean & Jean-Christophe, 2010; Matsubayashi et al., 2011).

Though ecological speciation is well-studied in one or two trophic level systems (Jiao et al., 2021; Matsubayashi et al. 2011), few studies have considered that shifts in biodiversity may also cascade across multitrophic systems. In this sequential divergence (Hood et al., 2015), also termed sequential speciation (Feder & Forbes, 2010) and cascading host-associated differentiation (Stireman et al., 2006), biodiversity at one trophic level ripples, affecting adaptation and divergence at other trophic levels (Feder & Forbes, 2010). Ultimately, cascading divergence can produce new niches for specialization in closely associated organisms (Figure 1). This may be especially relevant to biodiversity in multitrophic plant-insect-parasitoid systems. For example, in *Rhagoletis pomonella* fruit flies, multiple species of parasitoids have followed divergence of their host insects as they shifted from hawthorn to apple host trees (Feder & Forbes, 2010; Forbes et al., 2009; Hood et al., 2015). The internally feeding fruit fly’s plant shift resulted in two distinct fly races on two distinct host plants, with subsequent population divergence in each of three parasitoid species. Therefore, it is possible that diversification can beget further diversification when ecological divergence at one trophic level creates opportunities for parasitic associates to experience distinct selection pressures.

**Figure 1:**
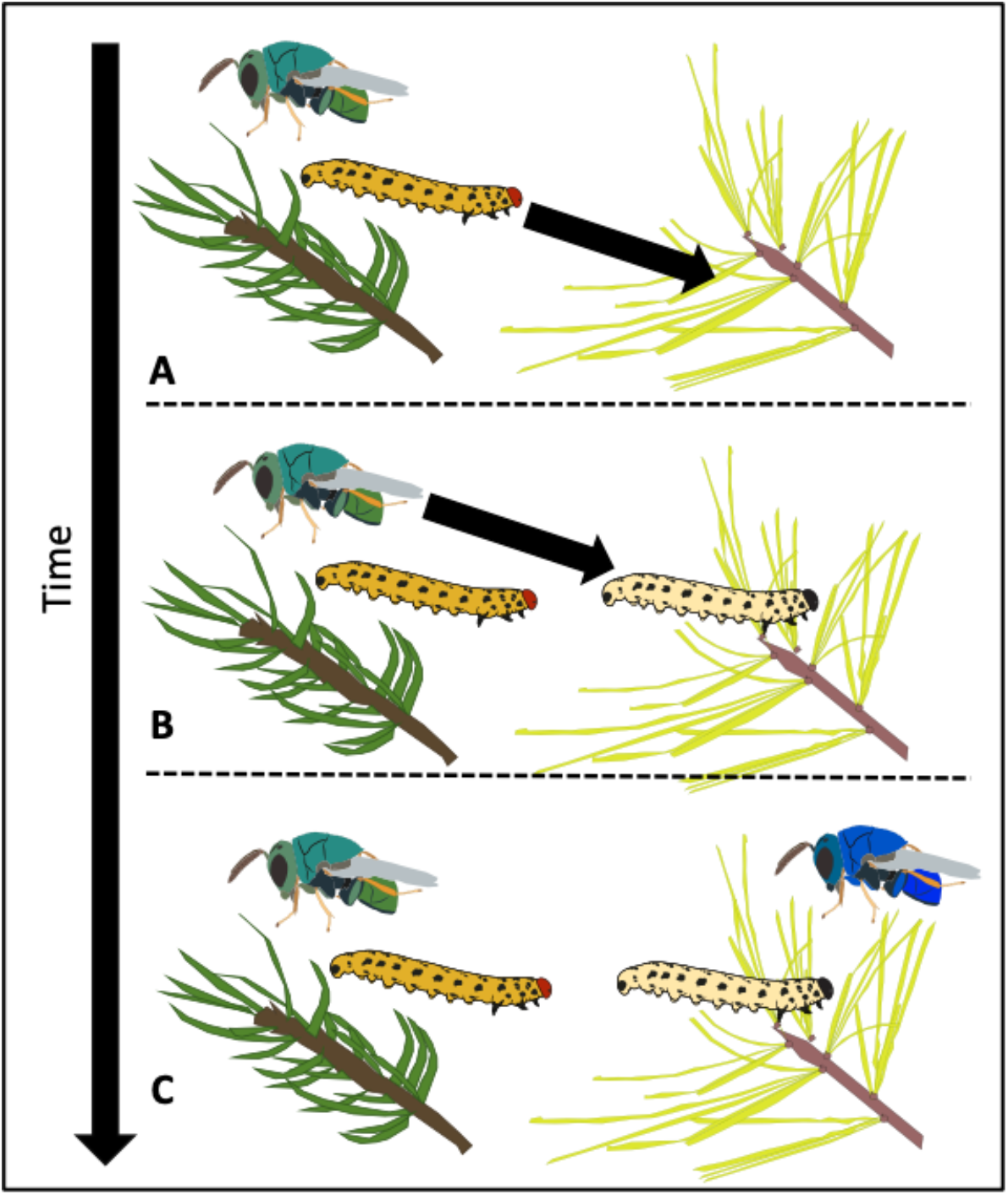
Sequential Divergence. A.) Plant-feeding insect shifts to new host plant. B.) Parasitoid wasp then shifts to newly diverged herbivorous insect. C). Divergence cascade produces two wasps on two host insects on two host plants. Alt text: Figure displaying process of sequential divergence. In panel A, a larval insect shifts onto a new pine species and undergoes divergence. In panel B, an associated parasitoid wasp undergoes a shift onto the newly diverged insect species. In panel C, two wasps, two insect hosts, and two pine trees are displayed.

Despite this striking example of sequential divergence in the fruit-*Rhagoletis*-parasitoid wasp system, research in this field is limited and demonstrates a gap in current biodiversity literature (Feder & Forbes, 2010; Forbes et al., 2009; Hood et al., 2015). As parasitoid wasps, especially those found in externally feeding herbivores, may harbor vast, unidentified species richness (Forbes et al., 2018), it is particularly relevant to determine the types of multitrophic systems in which sequential divergence drives parasitoid diversification. Currently, few other empirical studies of sequential divergence exist, and they have only positively identified sequential divergence in endophagous insects (e.g., parasitoids of gallmakers) (Baine et al., 2026; Blair & Abramson, 2008; Crespi & Abbot, 1999; Stireman et al., 2006; Thompson et al., 2022; Zhang et al., 2019).

No previous studies, to our knowledge, have supported sequential divergence as a driver of biodiversity in externally feeding insects and their parasitoids. However, only a handful of studies have explicitly investigated this scenario (see Baer et al., 2004; Bilodeau et al., 2013; Dickey & Medina, 2011; Lozier et al., 2008; Thompson et al., 2022). Furthermore, past studies may not have considered ideal systems for sequential divergence, as ecological factors (e.g., similar quality of host insects), natural history characteristics (e.g., multivoltinism, learned habitat preference), and short shared evolutionary histories (Bilodeau et al., 2013; Dickey & Medina, 2011; Simonato et al., 2012) may have slowed or inhibited sequential divergence. As plant-feeding insects comprise a quarter of all animal species, making ectophagous insects a considerable source of biodiversity (Bernays et al., 2009), additional research is needed to determine if externally feeding insects can ever serve as a focal point for cascading divergence. To address this idea, we require a system with externally feeding insect hosts that have strong plant-associated divergence, well-established natural histories, selection-driven habitat preference, and a historical plant-insect-parasitoid evolutionary relationship.

The externally feeding conifer sawfly genus *Neodiprion* (Smith, 1993) and its community of parasitoids present a viable system for further examining sequential divergence. *Neodiprion* sawflies exhibit close, lifelong associations with their *Pinus* hosts (Arnett, 1993; Benjamin, 1955), and both historical (Coppel & Benjamin, 1965; Knerer & Atwood, 1972; Knerer & Atwood, 1973) and recent research indicate that host shifts drive *Neodiprion* divergence (Linnen & Farrell, 2010). Furthermore, as a genus of pest insects, *Neodiprion* has unusually well-documented natural history (Arnett, 1993; Benjamin, 1955), including lists of its parasitoids (Forbes et al., 2018; Kephart, 2024).

One of these parasitoids, *Perilampus neodiprioni*, has a wide geographic and host range. It parasitizes at least 16 *Neodiprion* species throughout their North American ranges (personal observations; Tripp, 1962) and hyperparasitizes some primary parasitoids of *Neodiprion* (Yoo & Darling, 2024). Though specialized in its host range, *P. neodiprioni* harbors substantial morphological, ecological, and behavioral variation (Tripp, 1962), as well as possible genetic differentiation (Yoo & Darling, 2024), consistent with undescribed divergence. Furthermore, it is ideal for laboratory studies, because it is abundant and easily reared as by-catch of *Neodiprion* sawfly colonies (personal observations).

Recent research on body size across *Neodiprion* species (Glover et al., 2023) presents an opportunity to examine sequential divergence in the *Pinus-Neodiprion-P. neodiprioni* system (S. Figure 1). Glover et al. (2023) support body size as a “magic trait” in sympatric *Neodiprion* species with different host pine species. Magic traits, or those that simultaneously are adaptive and result in non-random mating, are one line of evidence for ecological speciation (Nosil et al., 2012; Servedio et al., 2011). In *Neodiprion,* body size acts as a magic trait because it responds to host-related divergent selection (different pine needle sizes create different selection pressures) and contributes to non-random mating (body size affects prezygotic isolation). On average, *N. lecontei* adults have significantly larger bodies than *N. pinetum* adults, a pattern closely associated with the distinct pine needle sizes of their different host plants (Glover et al., 2023). These habitat-driven differences in body sizes inhibit interbreeding of *N. lecontei* and *N. pinetum* throughout much of their overlapping geographic range.

Body size may be a candidate trait for driving cascading divergence across trophic levels. Past parasitoid research associated host identity with intraspecific body size sexual dimorphism (Bogdanović et al., 2009), but research has not assessed whether two host species’ body size divergence could create niches for parallel size divergence in parasitoids. By exploiting well-characterized pine-driven differences in *N. lecontei* and *N. pinetum’s* body sizes, we can assess whether *Neodiprion* sawfly host differences subsequently facilitate differences in *P. neodiprioni* body size. Such an analysis would provide novel insight into the multitrophic implications of magic trait-driven divergence. In particular, magic traits are widely accepted as one possible driver of ecological speciation (Nosil et al., 2012), and may be common and essential sources of divergence in habitat-associated organisms (Servedio et al., 2011). Since sequential divergence is understudied (Hood et al., 2015), it is unknown if magic can beget magic, leading to parallel divergence in a multitrophic cascade.

Here, we assess patterns of body size in the parasitoid of an externally feeding host – *Neodiprion* – and consider whether *Neodiprion*’s magic trait-based divergence facilitates similar patterns of divergence in its parasitoid, *Perilampus neodiprioni*. To assess the cascade of body size divergence, we photographed samples and measured body size across a wide geographic range of male and female *P. neodiprioni* reared from both *N. lecontei* and *N. pinetum* hosts. We used a generalized additive mixed model to analyze this body size data in relation to host species, host sex, host cocoon size, parasitoid sex, and latitude, with awareness of host species body size trends. We then used a mixed effects model with type III ANOVA and estimated marginal means tests to examine a subset of the data for host association and sexual dimorphism in parasitoids from the hybrid zone of two host species. We find that *P. neodiprioni* does exhibit host-specific divergence in body size that parallels differences found in their *Neodiprion* hosts, suggesting magic traits may ripple across trophic levels.

## Materials and Methods

### Sample Sorting

Parasitoids were obtained as by-catch from *Neodiprion* colonies collected as larvae between 2003 and 2024 by our laboratory and collaborators. *Neodiprion* sawflies were reared under conditions described by Bagley et al. (2017). Some collected sawfly colonies contained parasitized larvae, yielding parasitoids instead of adult sawflies upon eclosion. Eclosed parasitoids and their associated sawfly cocoon were stored individually, dry or in ethanol, at - 80°C.

Parasitoid specimens were identified to the species level using superfamily (Goulet & Huber, 1993) and genus keys (Townes, 1969; Wahl & Sharkey, 1993; Gibson et al., 1997; Cushman, 1940), as well as a key for parasitoids of *Neodiprion* (Finlayson, 1963) and a list of historical parasitoids of *Neodiprion* (Forbes et al., 2018). Species identities were corroborated with mitochondrial COI barcodes. Among these parasitoids, 2555 *Perilampus neodiprioni* specimens were obtained.

### Body Size Analysis

We selected 548 *P. neodiprioni* specimens representative of the parasitoid’s geographic and sawfly host species range. Samples were selected to maximize geographic variation and ensure good condition for measurement (i.e., we only considered specimens with intact bodies). For each sample, we placed the wasp in view on a Leica EZ4 stereo microscope. The sample was placed so that the head faced left, and the entire wasp and sawfly cocoon fit in the field of view. Specimens were photographed with a unique alphanumeric identifier to establish identity and a millimeter ruler to establish scale.

Body size measurements were obtained using ImageJ (Schneider et al., 2012). The in-frame ruler set the scale for each photograph. Total body length was obtained by summarizing anterior-to-posterior measurements of the head, thorax, and abdomen segments (S. Figure 2). In addition, we measured the cocoon size to act as a proxy for *Neodiprion* host body size, which represents resources available to the parasitoid prior to eclosion. Finally, we noted parasitoid sex and used cocoon morphology to identify the *Neodiprion* host’s sex (larger cocoons represent diploid females, smaller cocoons represent haploid males).

**Figure 2:**
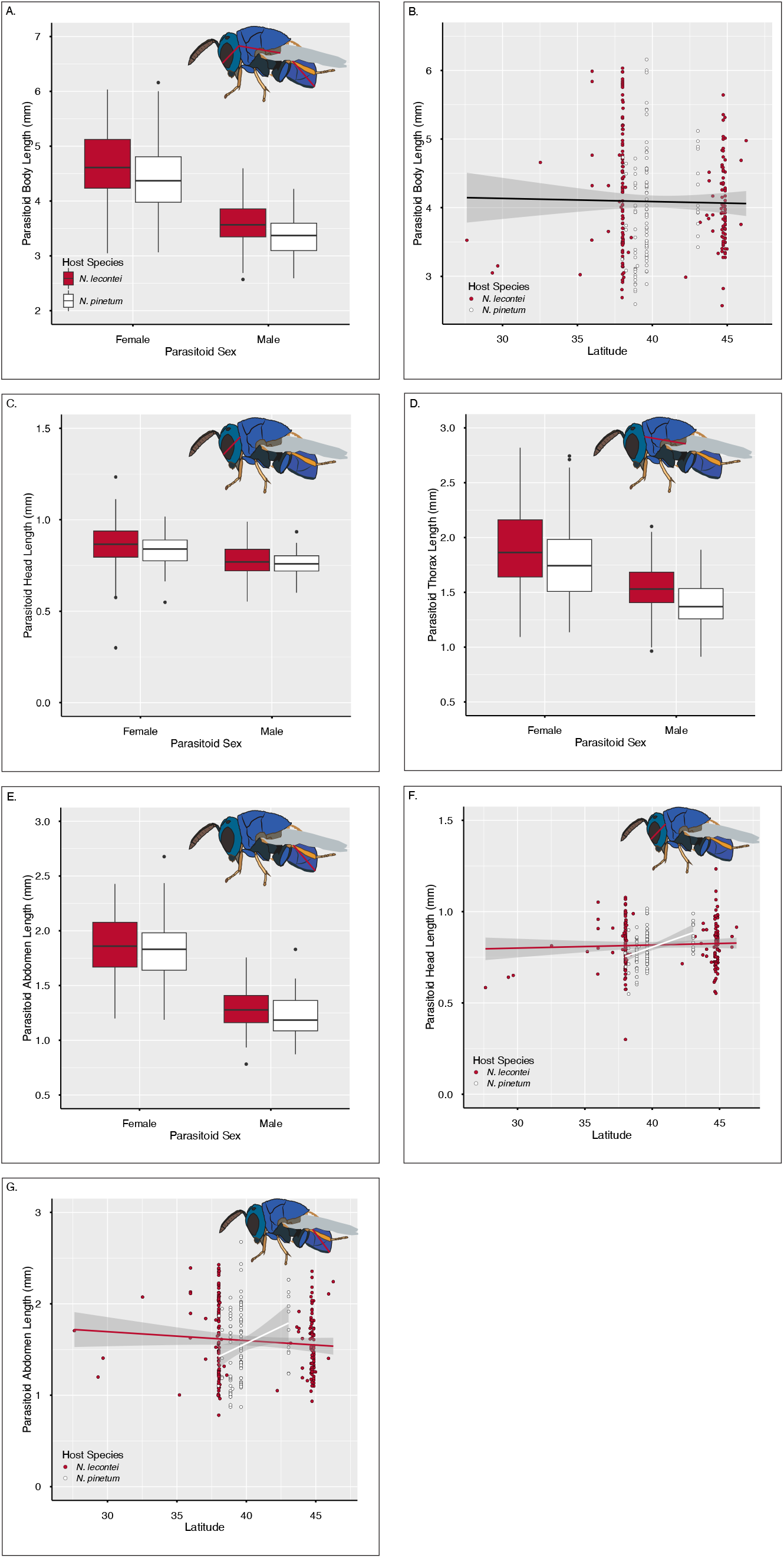
A.) Comparison of total body length of *P. neodiprioni* reared from *N. lecontei* and *N. pinetum* hosts. A Type III ANOVA and estimated marginal means contrast were used to evaluate body size variation between parasitoids reared from *N. lecontei* (n= 204) (red) and *N. pinetum* (n=103) (white) hosts, following a generalized additive mixed model. B.) Scatterplot with line of best fit depicting latitudinal cline of *P. neodiprioni* total body size length. C.-E.) Box plots visualizing size variation in each parasitoid body segment (C. Head, D. Thorax, E. Abdomen) for both male and female parasitoids from *N. lecontei* (red) and *N. pinetum* (white) hosts in the full geographic range analysis. F.) Scatterplot with lines of best fit depicting latitudinal clines of *P. neodiprioni’s* head size from *N. lecontei* (red) and *N. pinetum* (white) hosts. G.) Scatterplot with lines of best fit depicting latitudinal clines of *P. neodiprioni’s* abdomen size from *N. lecontei* (red) and *N. pinetum* (white) hosts. Alt text: Graphs displaying variation in parasitoid body size segments from *N. lecontei* and *N. pinetum* hosts. Subfigures labelled A to G, showing the described variation between male and female parasitoids and across latitudinal gradients.

To assess if and how body size varied among parasitoids, we focused on a subset of samples reared from sister sawfly species *N. lecontei* (n=204) and *N. pinetum* (n=103) (S. Figure 3). Using R Studio v4.4.2 (R Studio Team, 2020), we proposed a mixed model with lme4 v2.0-6 (Bates et al., 2015) to determine how continuous parasitoid body size varies with host species (categorical), host sex (categorical), host cocoon size (continuous), parasitoid sex (categorical), and latitude (continuous), as well as the paired interactions of these variables. While host species identity is our primary variable of interest, we included the additional explanatory variables in our initial model, as they are likely to impact body size (Glover et al., 2023). Host colony identity was included as a random effect, as parasitoids from the same host colony could possibly be siblings and have increased similarity.

**Figure 3:**
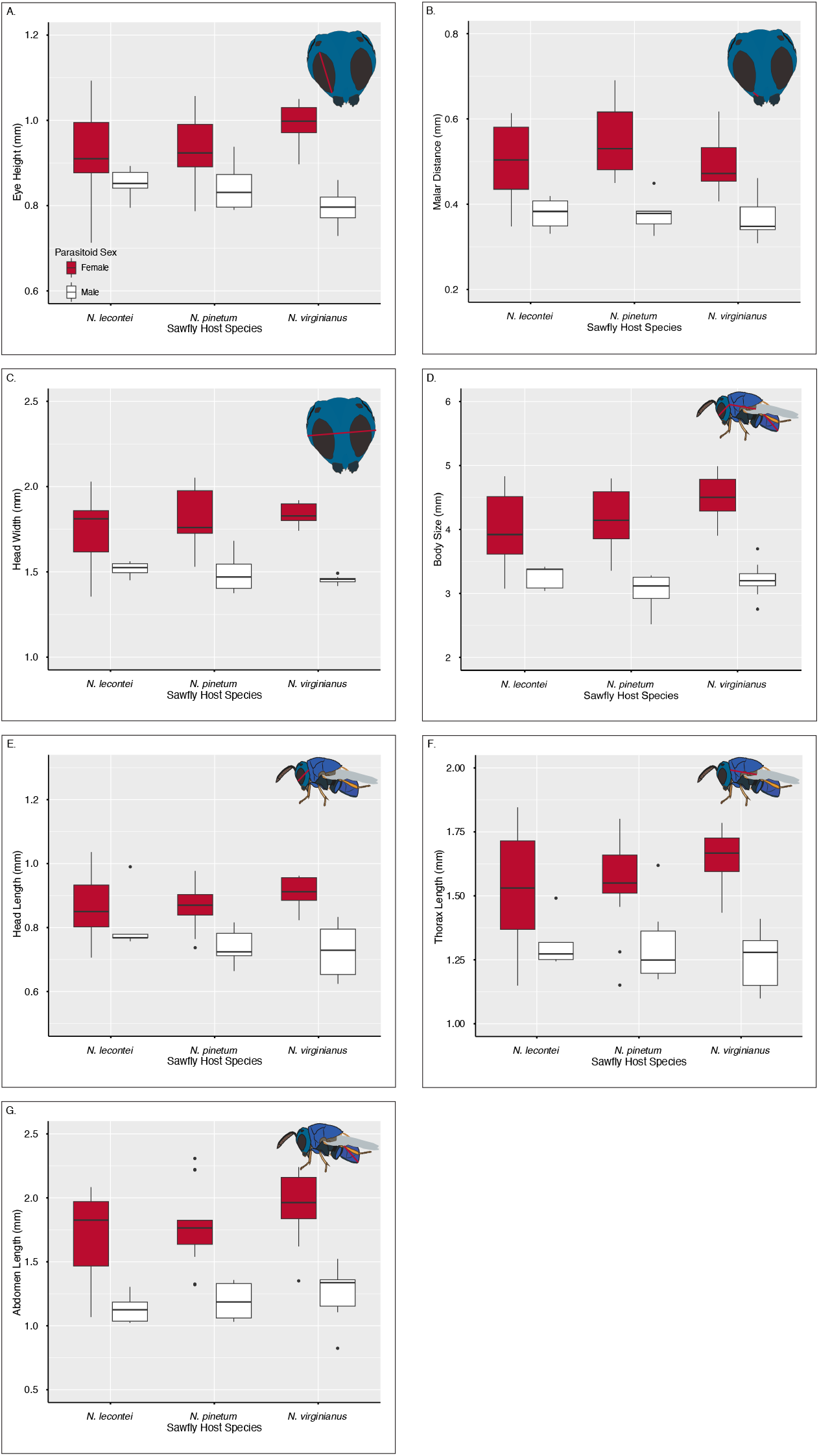
A.) Box plots visualizing size variation in limited-geography analysis of *P. neodiprioni* from different host species. Each plot portrays a parasitoid trait of interest (A. Eye Height, B. Malar Distance, C. Head Width, D. Total Body Size, E. Head Length, F. Thorax Length, G. Abdomen Length) for both male (white) and female (red) parasitoids from *N. lecontei*, *N. pinetum,* and *N. virginianus* hosts. Alt text: Graphs displaying host-associated and sex-based size variation in facial features and body segments of parasitoids. Subplots are labelled A to G, with each plot depicted one trait.

After creating our initial model, interaction terms that were not significant (p<0.05) or marginally significant (p<0.1) were removed to reduce overfitting. Then, we checked model assumptions (linearity, normality, homogeneity, independence, normality of random effects) with the DHARMa v0.4.7 (Hartig, 2024) and performance v0.17.1 (Lüdecke et al., 2021) packages. To address minor assumption deviations, the model was converted to a generalized additive model in mgcv v1.7-10 (Wood & Wood, 2015), with the colony variable as a penalized regression term with random effect basis. After reassessing assumptions in mgcv v1.7-10 (Wood & Wood, 2015), we further explored results via a type III analysis of variance (ANOVA) with mgcv v1.7-10 (Wood & Wood, 2015) and estimated marginal means with emmeans v2.0.4 (Lenth, 2025). Results were compared to *Neodiprion* body size analyses in Glover et al. (2023) to facilitate interpretation.

After identifying body size variation in *P. neodiprioni*, we sought to determine, more specifically, which body segments exhibit host-associated variation. To this end, we repeated model evaluation for each individual body segment: head, thorax, and abdomen. For the thorax and abdomen analyses, the originally proposed mixed effects model was created, nonsignificant interaction terms were removed, the model was converted to a generalized additive model to smooth variables causing minor assumption deviations, and assumptions were re-checked with visual plots and statistical outputs in mgcv v1.7-10 (Wood & Wood, 2015). Follow-up type III ANOVAs and estimated marginal means analyses were performed.

To address singularity in the head model caused by the colony random effect, a multiple linear regression model was evaluated instead. The same previously proposed fixed effects were analyzed, with non-significant interaction terms removed and model assumptions assessed. We then performed a type III ANOVA and estimated marginal means analyses.

### Sexual Dimorphism Data Collection

After noting that host-associated size variation occurred in the abdomen, we sought to understand if size variation could impact mating. As dead specimens prohibited live mating assays, we considered if other size-based features, which typically exhibit sexual dimorphism, have host-driven variation. To consider host effects on size-based, sexually dimorphic characters in *P. neodiprioni*, we examined 70 specimens, spread across 3 sympatric host species: *N. lecontei, N. pinetum,* and *N. virginianus.* Parasitoids from *N. virginianus* were added to this analysis to act as an outgroup, as all samples were obtained from a short latitudinal transect from Lexington, Kentucky to southern Ohio. This controlled geographic range represents the hybrid zone of the two sister species (*N. lecontei* and *N. pinetum*), but not *N. virginianus*.

Samples were photographed laterally and body segments were measured as described in the body size analysis. We also obtained a ventral photograph. The eye height (Yoo & Darling, 2024), malar length (Mathou et al., 2023; Yoo & Darling, 2024), and head width (King et al., 2018) were measured from the lateral and ventral images (S. Figure 2). The ventral photograph was also used to identify the sample’s sex, based on genitalia.

We analyzed size variation in each character of interest, independently: eye height, malar distance, head width, total body size, head size, thorax size, and abdomen size. For each character, we used a mixed effects model (lme4 v2.0-6 (Bates et al., 2015)) to determine whether the average size of each trait varies across parasitoid sex and *Neodiprion* host species. We included the explanatory variables’ interaction term and host colony as a random effect. Model assumptions were checked with DHARMa v0.4.7 (Hartig, 2024) and performance v0.17.1 (Lüdecke et al., 2021). Following model evaluation, the colony random effect was removed from the eye height analysis due to singularity, and the model was run as a multiple regression model. In the head width analysis, the host species by parasitoid sex interaction term was weighted in nlme v3.1-152 (Pinheiro et al., 2025) to allow different residual variance that addresses heteroskedasticity. In the abdomen model, the non-significant interaction term was removed to reduce overfitting. To further explore results, we performed a type III ANOVA with car v3.1-0 (Fox & Weisberg, 2019) and estimated marginal means analyses with emmeans v2.0.4 (Lenth, 2025) for each trait.

## Results

### Total Body Size Results

Our generalized additive mixed model of total parasitoid body size has a strong fit (R^2^_adj_=0.776, Deviance Explained = 82%, Estimated df = 61.34), and it indicates several key relationships (Table 1). First, total *P. neodiprioni* body size varies marginally significantly with *Neodiprion* host species identity (p=0.055). This association occurs even with the inclusion of cocoon size, which accounts for variation in resources provided by individual hosts. *P. neodiprioni* reared from *N. lecontei* tend to be larger, on average, than *P. neodiprioni* reared from *N. pinetum* (S. Table 1) (Figure 2A). Based on the non-significance of the host by parasitoid sex interaction, this occurs regardless of parasitoid sex; both male and female parasitoids from *N. lecontei* are larger on average than counterparts from *N. pinetum*.

**Table 1:**
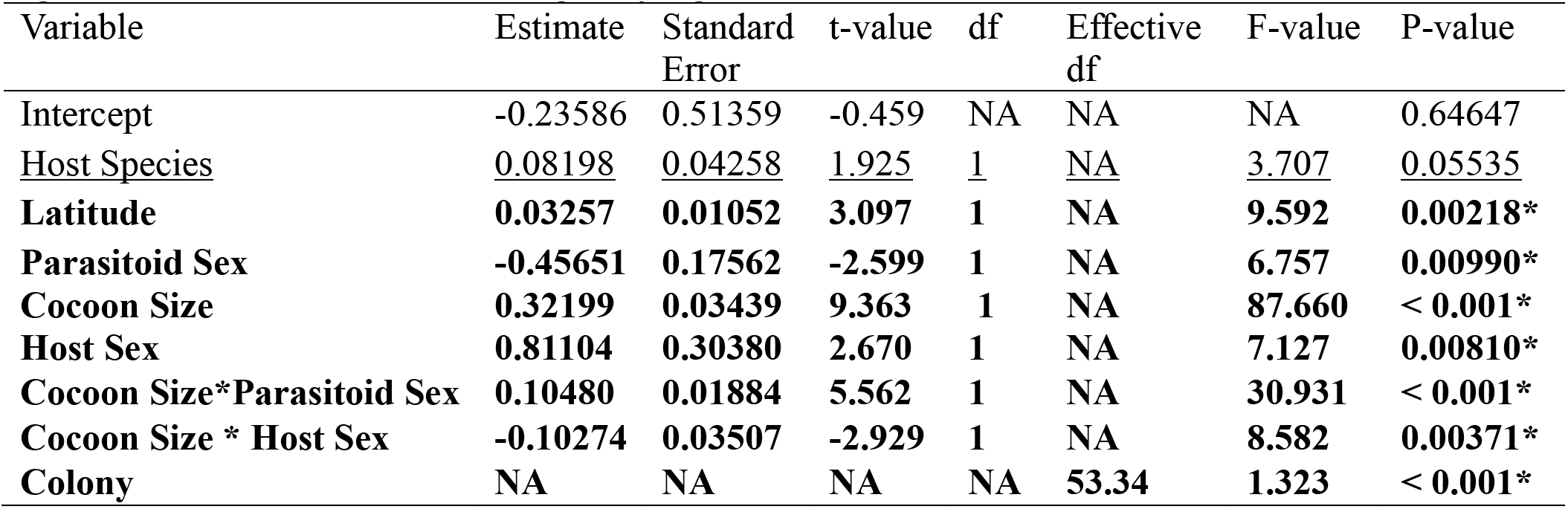
Statistical outputs of generalized additive mixed model (R^2^_adj_=0.776, Deviance Explained = 82%, n=307) and associated type III ANOVA for full-range body analysis (Parasitoid Body Size ∼ Host Species + Latitude+ Host Sex + Parasitoid Sex + Cocoon Size + Cocoon Size*Parasitoid Sex + Cocoon Size * Host Sex + smooth(Colony, random effect)). Significant terms are bolded and marginally significant terms are underlined.

| Variable | Estimate | Standard Error | t-value | df | Effective df | F-value | P-value |
|---|---|---|---|---|---|---|---|
| Intercept | -0.23586 | 0.51359 | -0.459 | NA | NA | NA | 0.64647 |
| Host Species | 0.08198 | 0.04258 | 1.925 | 1 | NA | 3.707 | 0.05535 |
| **Latitude** | **0.03257** | **0.01052** | **3.097** | **1** | **NA** | **9.592** | **0.00218\*** |
| **Parasitoid Sex** | **-0.45651** | **0.17562** | **-2.599** | **1** | **NA** | **6.757** | **0.00990\*** |
| **Cocoon Size** | **0.32199** | **0.03439** | **9.363** | **1** | **NA** | **87.660** | **< 0.001\*** |
| **Host Sex** | **0.81104** | **0.30380** | **2.670** | **1** | **NA** | **7.127** | **0.00810\*** |
| **Cocoon Size\*Parasitoid Sex** | **0.10480** | **0.01884** | **5.562** | **1** | **NA** | **30.931** | **< 0.001\*** |
| **Cocoon Size \* Host Sex** | **-0.10274** | **0.03507** | **-2.929** | **1** | **NA** | **8.582** | **0.00371\*** |
| **Colony** | **NA** | **NA** | **NA** | **NA** | **53.34** | **1.323** | **< 0.001\*** |

Latitude has a significant association with *P. neodiprioni* body size. Overall, as latitude increases, *P. neodiprioni* body size has minor increases (Table 1) (Figure 2B). Model selection indicated that latitude does not significantly interact with other variables.

*P. neodiprioni* body size also has a significant relationship with host cocoon size (Table 1). As cocoon size increases, *P. neodiprioni* body size also increases. In addition, the cocoon size and parasitoid sex interaction is significant (Table 1), where female parasitoids tend to have a greater rate of increase in body size with increases in *Neodiprion* cocoon size than their male counterparts (S. Table 2) (S. Figure 4), and female parasitoids are larger overall than male parasitoids (S. Table 1). Furthermore, the significant interaction of host cocoon size and host sex indicates that parasitoids from male hosts have a greater rate of increase in average body size with increases in host cocoon size than parasitoids from female hosts (S. Table 2). Therefore, increased cocoon size is associated with increased parasitoid body size, but the rate of this increase varies depending upon parasitoid sex and host sex.

Finally, the smoothed colony random effect has a significant association with parasitoid body size, indicating that, range-wide, parasitoids from the same host colony are more similar to one another in size than would be expected by random chance alone (Table 1).

### Body Segment Results

Upon repeating model selection, and model, ANOVA, and estimated marginal means analyses for each insect body segment independently, all three body segments converged, but those models varied in strength and significant terms (S. Table 3, S. Table 4, S. Table 5).

The multiple regression model of head size variation is significant but weak (R^2^_adj_=0.311, df= 299, p < 0.001) (S. Table 3) (Figure 2C). Both host species and latitude are significant, as well as their interaction. As latitude increases, the head size of parasitoids from *N. pinetum* hosts tends to increase at a greater rate than the increase in head size of parasitoids from *N. lecontei* hosts (Table 2). In addition, the cocoon size and host sex variables, as well as their interaction, are significant. Here, increases in cocoon size are associated with increases in parasitoid head size, but parasitoids from male hosts have a greater rate of increase with cocoon size compared to counterparts from female hosts (Table 2). Finally, parasitoid sex has a significant association with head size, where female parasitoids have larger heads on average than males (Table 2).

**Table 2:** Paired comparisons from estimated marginal means analyses of head, thorax, and abdomen models in full geographic range analysis. For interpretative clarity, variables that are included in a significant interaction term are only presented in the interaction term’s estimated marginal means analysis, not each individual variable’s estimated marginal means analysis. Significant terms are bolded and marginally significant terms are underlined.

| Model | Variable | Contrast | Estimate | Standard Error | df | t-ratio | P-value |
|---|---|---|---|---|---|---|---|
| **Head** | **Parasitoid Sex** | **Female - Male** | **0.0794** | **0.0111** | **299** | **7.138** | **< 0.001\*** |
|  | **Latitude\*Host Species** | ***N. lecontei*** **–** ***N. pinetum*** | **-0.0165** | **0.00725** | **299** | **-2.273** | **0.0237\*** |
|  | **Cocoon Size\*Host Sex** | **Female - Male** | **-0.0321** | **0.0154** | **299** | **-2.085** | **0.0380** |
| **Thorax** | Host Species | *N. lecontei* – *N. pinetum* | 0.0884 | 0.0513 | 251 | 1.723 | 0.0861 |
|  | Latitude* Parasitoid Sex | Female - Male | -0.0215 | 0.0111 | 251 | -1.931 | 0.0546 |
|  | **Cocoon Size\* Host Sex** | **Female - Male** | **-0.103** | **0.0442** | **251** | **-2.336** | **0.0203** |
|  | **Latitude\*Host Sex** | **Female - Male** | **0.047** | **0.0167** | **251** | **2.814** | **0.0053** |
| **Abdomen** | **Host Sex** | **Female - Male** | **0.0137** | **0.0381** | **257** | **0.359** | **0.7196** |
|  | **Cocoon Size\* Parasitoid Sex** | **Female - Male** | **0.0896** | **0.0175** | **257** | **5.113** | **< 0.001\*** |
|  | **Latitude\*Host Species** | ***N. lecontei*** **–** ***N. pinetum*** | **-0.0511** | **0.0221** | **257** | **-2.310** | **0.0217** |

The generalized additive mixed model of thorax size has moderate strength (R^2^_adj_=0.604) (S. Table 4). It indicates a marginally significant association of host species with thorax size, where parasitoids from *N. lecontei* hosts have larger thoraxes on average than parasitoids from *N. pinetum* hosts (Table 2) (Figure 2D). The cocoon size by host sex interaction is also significant, where parasitoids from male hosts have a greater rate of thorax size increase with cocoon size increase (Table 2). In addition, though both cocoon size and latitude have positive associations with thorax size, cocoon size’s association becomes less positive as latitude increases (S. Table 4). Latitude also interacts significantly with host sex and marginally significantly with parasitoid sex (Table 2). Parasitoids from female hosts have a greater rate of increase in thorax size with increased latitude than male counterparts (Table 2). Female parasitoids exhibit a small decrease in thorax size with increases in latitude (estimate = -0.0065 +/-0.00824), while male parasitoids have a pattern of increased thorax size with increases in latitude (estimate = 0.0150 +/-0.00989) (Table 2). Finally, the significant interaction of cocoon size and host sex indicates that parasitoids from female hosts have a less positive rate of increase in thorax size with cocoon size than parasitoids from male hosts (Table 2).

Finally, the model of abdomen size has a strong fit (R^2^_adj_=0.785). Abdomen size is significantly associated with *Neodiprion* host and the interaction of host species and latitude (S. Table 5). The abdomen size of parasitoids from *N. lecontei* decreases with increased latitude compared to the increase in parasitoids from *N. pinetum* (Table 2). This, combined with the non-significant estimated marginal means of host species alone (p=0.9359) (Figure 2E), indicates that parasitoids from *N. lecontei* and *N. pinetum* maintain distinct abdomen sizes throughout their latitudinal range, but parasitoids from one host species do not simply have larger abdomens. There is also a significant interaction of cocoon size and parasitoid sex, where abdomen size increases on average with increases in cocoon size (S. Table 5), but female parasitoids have a greater rate of abdomen size increase than males (Table 2). Finally, host sex has a significant association with abdomen size, where parasitoids from female hosts are larger on average than parasitoids from male hosts (Table 2).

### Sexual Dimorphism Results

Analyses of three morphological features of *P. neodiprioni*, eye height, malar distance, and head width, show significant size-based variation in association with parasitoid sex, which sometimes significantly interacts with host species (Table 3) (S. Table 6). Eye height, head width, and malar distance are larger on average among female parasitoids than male parasitoids (Figure 3). When considering sex-related variation of eye height between host species, there is a marginally significant pattern, where females from *N. lecontei* hosts have smaller eye heights on average than females from *N. virginianus* hosts (Table 3). Female parasitoids from *N. pinetum* are not significantly different in eye height from *N. lecontei* or *N. virginianus*. For sex-related variation of head width between host species, male parasitoids from *N. virginianus* exhibit smaller head widths on average than males from *N. lecontei* hosts (Table 3). Malar distance does not exhibit a significant parasitoid sex by host species interaction.

**Table 3:** Estimated marginal means analyses of explanatory variables in eye height, malar distance, and head width models in the limited-geography range. For interaction terms of eye height and head width models, pairwise comparison of the parasitoids’ facial traits considered sawfly host species within each parasitoid sex grouping, and all comparisons used a Tukey adjustment. For interpretative clarity, variables that are included in a significant interaction term are only presented in the interaction term’s estimated marginal means analysis, not each individual variable’s estimated marginal means analysis. The estimated marginal means analysis of malar distance considered the two explanatory variables seperately, due to the non-signfiicant interaction term. Significant terms are bolded and marginally significant terms are underlined.

| Model | Variable | Group | Contrast | Estimate | Standard Error | df | t-ratio | P-value |
|---|---|---|---|---|---|---|---|---|
| Eye Height | Parasitoid Sex* Host Species | Female | *N. lecontei* – *N. pinetum* | -0.01052 | 0.0265 | 58 | -0.398 | 0.9166 |
|  |  | Female | *N. lecontei* – *N. virginianus* | -0.06739 | 0.0305 | 58 | -2.211 | 0.0777 |
|  |  | Female | *N. pinetum* - *N. virginianus* | -0.05687 | 0.0324 | 58 | -1.753 | 0.1945 |
|  |  | Male | *N. lecontei* – *N. pinetum* | 0.00813 | 0.0460 | 58 | 0.177 | 0.9829 |
|  |  | Male | *N. lecontei* – *N. virginianus* | 0.05720 | 0.0416 | 58 | 1.375 | 0.3603 |
|  |  | Male | *N. pinetum* - *N. virginianus* | 0.04907 | 0.0392 | 58 | 1.251 | 0.4282 |
| Malar Distance | **Parasitoid Sex** | **NA** | **Female - Male** | **0.133** | **0.0211** | **52** | **6.286** | **< 0.001\*** |
|  | Host Species | NA | *N. lecontei* – *N. pinetum* | -0.03063 | 0.0271 | 28.1 | -1.132 | 0.5026 |
|  | Host Species | NA | *N. lecontei* – *N. virginianus* | -0.00172 | 0.0255 | 33.6 | -0.067 | 0.9975 |
|  | Host Species | NA | *N. pinetum* - *N. virginianus* | 0.02892 | 0.0167 | 28.0 | 1.034 | 0.5620 |
| Head Width | Parasitoid Sex* Host Species | Female | *N. lecontei* – *N. pinetum* | -0.0537 | 0.0612 | 21 | -0.877 | 0.6603 |
|  |  | Female | *N. lecontei* – *N. virginianus* | -0.0774 | 0.0453 | 21 | -1.708 | 0.2258 |
|  |  | Female | *N. pinetum* - *N. virginianus* | -0.0238 | 0.0502 | 21 | -0.474 | 0.8842 |
|  |  | Male | *N. lecontei* – *N. pinetum* | 0.0245 | 0.0492 | 21 | 0.498 | 0.8728 |
|  |  | **Male** | ***N. lecontei* – *N. virginianus*** | **0.0616** | **0.0199** | **21** | **3.093** | **0.0146\*** |
|  |  | Male | *N. pinetum* - *N. virginianus* | 0.0370 | 0.0461 | 21 | 0.804 | 0.7045 |

In the limited-geography analysis of body size via a mixed model, parasitoid sex varies significantly with body size, where females are larger on average than males (Figure 3). The interaction of the parasitoid sex and host species terms had marginal significance, warranting post-hoc estimated marginal means analysis (S. Table 7). Here, female parasitoids from *N. lecontei* hosts are smaller on average than female parasitoids from *N. virginianus*, though neither have significant differences when compared to female parasitoids from *N. pinetum* (S. Table 8).

When each body segment is examined individually in the limited geography dataset, similar patterns persist. For all three segments, head, thorax, and abdomen, parasitoid sex is significantly associated with size, and females are larger on average than males (S. Table 8) (Figure 3). In addition, both the head and thorax models show significant variation of size with the parasitoid sex by host species interaction. Head size is significantly different between males and females when samples come from *N. pinetum* and *N. virginianus* hosts, but not from *N. lecontei* hosts (S. Table 8). In the thorax model, females are significantly larger than males among samples derived from *N. lecontei* and *N. virginianus*, but not *N. pinetum*. In addition, there is a marginally significant pattern where thorax size in females from *N. virginianus* is larger on average than thorax size in females from *N. lecontei* (S. Table 8). In contrast, the parasitoid sex by host species interaction was dropped from the abdomen mixed model due to non-significance, but host species shows marginal significance in its association with abdomen size (S. Table 8). Parasitoids from *N. lecontei* have smaller abdomens on average than parasitoids from *N. virginianus* (Figure 3).

## Discussion

*P. neodiprioni* exhibits initial evidence of host-associated divergence in body size, which is broadly indicative of ecological speciation. As differentiation in *P. neodiprioni* body size parallels the magic body size differences of its divergent *Neodiprion* hosts (Glover et al., 2023), this host-associated variation suggests that sequential divergence, or cascading ecological divergence, may occur in *P. neodiprioni.* Host plays a role in body size variation across body segments, but the host-associated body size variation is particularly notable in the abdomen region, raising questions of mechanical isolation. In addition, in a geographically-constrained dataset that includes the *N. lecontei* and *N. pinetum* hybrid zone, size-based facial features and body segments demonstrate both sexual dimorphism and parasitoid sex by host species interactions. Parasitoids from the non-sister *N. virginianus* host are often more different from parasitoids of the sister *N. lecontei* and *N. pinetum* hosts than parasitoids from the sister host species are from each other. Together, this evidence gives the first indication that sequential divergence may be possible in this system and among parasitoids of externally feeding insects in general.

### Host-Associated Body Size Divergence

Morphological divergence is well-established as one line of evidence for ecological speciation (Nosil, 2012; Langerhans et al., 2007), and body size has supported divergence in both plant-eating insects (Glover et al., 2023) and host-associated parasitoids (Morehead et al., 2001). Host-associated morphological variation alone does not indicate that a host shift drives speciation (Forbes et al., 2017; Schluter, 2009), but morphological analyses can support ecological speciation when combined with genetic, ecological, behavioral, mating, or fitness-based analyses (Glover et al., 2023; Forbes et al., 2017; Nosil, 2012; Langerhans et al., 2007). In fact, morphology is key to several tests of ecological divergence, including magic trait-based approaches (Nosil, 2012). When testing for magic traits, one assesses if a particular trait is under divergent natural selection across environments while also contributing to non-random, assortative mating (Nosil, 2012).

In *Neodiprion*, ecological speciation is supported by body size as a magic trait in the sister species of *N. lecontei* and *N. pinetum* (Glover et al., 2023). Here, body size is under natural selection (different body sizes are better suited to different pine tree species) and contributes to non-random mating. As body size is a magic trait in *Neodiprion* and hypothesized as a common magic trait in specialized parasitoids (Glover et al., 2023), it was a useful first characteristic to test for host-associated divergence in *P. neodiprioni*.

*P. neodiprioni* displays marginally significant morphological divergence of body size associated with *Neodiprion* host species (Figure 2A). As different host insects can act as different environments for parasitoids (Hood et al., 2015; Aldrich & Zhang, 2002; Morehead et al., 2001), *P. neodiprioni* thus displays morphological divergence across environments. This initial evidence of host-associated morphological divergence indicates that ecological speciation could occur among these parasitoids. If *P. neodiprioni* body size is under divergent natural selection and impacts mating, it would confirm ecological speciation (Glover et al., 2023) and support sequential divergence.

Notably, the host-associated body size divergence of *P. neodiprioni* parallels the body size divergence of their *Neodiprion* hosts. Just as *N. lecontei* are larger than *N. pinetum* (Glover et al., 2023), *P. neodiprioni* reared from *N. lecontei* are larger than parasitoids from *N. pinetum*. This parallel morphological divergence is particularly important because parallel ecological divergence is suggestive of sequential divergence.

The parallel pattern of environment-associated differences in *Neodiprion* and *P. neodiprioni,* in that the same trait (body size) has similar trends (smaller *N. pinetum* and *N. pinetum* associates), presents initial support for sequential divergence in this system. If host-associated ecological speciation occurs in *P. neodiprioni,* it would indicate a divergence cascade from pine-associated *Neodiprion* to *Neodiprion*-associated *P. neodiprioni.* In particular, if *P. neodiprioni* size drives reproductive isolation, size would act as a magic trait in *P. neodiprioni*, just as it does in *Neodiprion*. This would suggest that magic traits can beget magic traits. However, morphological divergence alone does not indicate a magic trait or an ecological speciation cascade, requiring further investigation.

### Morphological Evidence of Possible Reproductive Isolation

Establishing a magic trait requires evidence of both divergent natural selection and effects on reproductive isolation (Nosil, 2012). This task is difficult using morphology alone, and we were unable to perform mating assays due to the preserved nature of the specimens. However, we considered possible means of size-driven reproductive isolation, crucial to magic phenomena, using phenotypic data.

First, we considered which body segments of *P. neodiprioni* display host-associated size variation. Though the head model shows a significant association with host species, the heads of parasitoids were slightly larger in samples from *N. pinetum* hosts, and estimated marginal means indicate this is actually not significant (p=0.9029). However, the significant host species by latitude interaction suggests that parasitoids from distinct hosts maintain different head size trends throughout their range (Table 2). The thorax model shows that parasitoids from *N. lecontei* hosts tend to have larger thoraxes, just as *N. lecontei* tend to be larger than *N. pinetum,* but this is a marginally significant association. Finally, the abdominal region of the parasitoid wasps demonstrates significant variation with host species over the latitudinal gradient that parallels a key pattern in *Neodiprion* hosts: parasitoids from *N. lecontei* and *N. pinetum* have distinct average abdomen sizes throughout the majority of their range (Figure 3). An intersection in average abdomen size occurs once, just north of the host hybrid zone.

*P. neodiprioni’s* largely distinct, host-associated abdomen sizes, except in close proximity to their hosts’ hybrid zone, provides an initial indication of possible parallel isolation patterns in the host and parasitoid species. It is notable that this occurs in the abdomen, as the abdomen is the insect body segment containing genitalia and reproductive structures (Imms, 1925; Snodgrass, 1931). Genitalia-related mechanical incompatibilities play a key role in prezygotic isolation among insect species (Barnard et al., 2017; Nagata et al., 2009; Tanaka et al., 2018). In addition, studies in insects (Glover et al., 2023; Okuzaki et al., 2010; Villa et al., 2019; Weissman et al., 2008), as well as research throughout animal groups (Conde-Padín et al., 2008; Ólafsdóttir et al., 2006; Tanabe & Sota, 2008), have demonstrated that adult body size differences in sympatric groups can strongly reduce interspecific reproduction. Therefore, clear body size differences in *P. neodiprioni,* especially abdominal size differences, could play a role in reducing reproduction between parasitoids reared from distinct *Neodiprion* hosts.

To further examine mechanisms of possible reproductive isolation, we considered commonly sexually dimorphic, size-based features of insects (Bogdanovic et al., 2009) that may potentially impact mate choice. These features were located in the head region: eye height (Yoo et al., 2024), malar distance (Mathou et al., 2023; Yoo et al., 2024), and head width (King et al., 2018). All traits were considered in male and female parasitoids from three sympatric host species, *N. lecontei, N. pinetum,* and *N. virginianus*, along a Central Kentucky through Southern Ohio transect. Here, the species occur in sympatry, even co-occurring at single sites (personal observation). This is also the only region where the sister species *N. lecontei* and *N. pinetum* have compatible sizes and have been observed to hybridize in the wild (Bendall et al., 2017). Body size analyses were also repeated on this smaller dataset to establish how parasitoid size varies in the host hybrid zone.

From these geographically-constrained facial feature and size analyses, *P. neodiprioni* shows consistent evidence of sexual dimorphism (larger female traits), which sometimes varies across host species (Figure 3). Female parasitoids of *N. virginianus* diverged from female parasitoids of *N. lecontei* in eye height, body size, and thorax size; male parasitoids of *N. virginianus* have shorter average head widths than males from *N. lecontei*; and parasitoids, regardless of sex, from *N. virginianus* have larger abdomens on average than parasitoids from *N. lecontei*.

In all cases, host comparisons within parasitoid sex groupings only demonstrated host-related effects between *N. virginianus* parasitoids and parasitoids from *N. lecontei*, but never between parasitoids from *N. lecontei* and *N. pinetum*. *N. virginianus* is more distantly related to *N. lecontei* and *N. pinetum* than these sister species are to each other (Herrig et al., 2024). Despite all three species occurring in sympatry, *N. virginianus* has not been observed to hybridize with the other two species in the wild. Therefore, our results provide an initial suggestion that host effects in sexually dimorphic parasitoid features may vary in parallel with divergence of host species. Just as *N. lecontei* and *N. pinetum* are more closely related and less severely reproductively isolated, the parasitoids from *N. virginianus* show initial evidence of being more significantly morphologically diverged from parasitoids from *N. lecontei* than the parasitoids of *N. lecontei* and *N. pinetum* are from one another. This suggests the reduced isolation of their sawfly hosts may allow greater opportunities for *P. neodiprioni* associated with *N. pinetum* and *N. lecontei* in this region to exchange genes, reducing their degree of morphological differentiation.

Parasitoids from different *Neodiprion* hosts in the Kentucky-Ohio range likely experience different ecological conditions to drive these host-associated sexual dimorphism differences. This phenomenon has been recorded in other parasitic wasps (Mathou et al., 2023) and would support a degree of reproductive isolation between parasitoids from different host environments (Martin & Mendelson, 2016; Mendelson, 2007). Such reproductive isolation is crucial to identifying ecological speciation. Thus, host-variable sexual dimorphism in this system prompts further investigation to assess if size-based traits truly inhibit reproduction and thereby facilitate sequential divergence in *P. neodiprioni*.

After identifying trends that indicate divergence of parasitoids from *N. lecontei* and *N. pinetum* throughout their range, but not in the area that aligns with their hosts’ hybrid zone, we considered that the *P. neodiprioni* may exhibit the same hybrid zone as their hosts. To address this possibility, we performed an additional analysis of parasitoids from *N. lecontei* and *N. pinetum* in a small northernly range, to compare host-associated divergence within and without the host hybrid zone. To do this, we first performed a type-III ANOVA on total body size variation of 35 *P. neodiprioni* samples from 12 *N. lecontei* colonies at the same site. This assessed whether parasitoids from the same host colony are more similar to one another than one would expect from random sampling within a small region. Results do not indicate a significant difference in size across colonies (p=0.07338), though future work must more comprehensively assess similarity within and between colonies.

After establishing that, within a small region, *P. neodiprioni* from the same colony are not more similar than predicted by random chance alone, we selected 13 *N. pinetum* and 16 *N. lecontei* from a northern latitudinal range (42.2287-44.658167). We performed a multiple regression of body size (R^2^_adj_=0.536, df=26, p<0.001), as predicted by host species and sex. We excluded the colony random effect due to the limited colony availability. Results indicate a significant association of parasitoid sex (p<0.001) and host species (p=0.0117), but not their interaction. This suggests that, unlike in the Kentucky-Ohio analysis, *P. neodiprioni* from *N. lecontei* and *N. pinetum* sister host species are morphologically divergent in northern regions. This contrast further supports that *P. neodiprioni* may be divergent in much of their range, but morphologically similar in their hosts’ hybrid zone.

### Body Size Divergence With Two Host Traits: Size and Sex

Beyond host species identity, two host traits, cocoon size and host sex, have strong associations with parasitoid body size in the full-range dataset. As host cocoon size increases, parasitoid body size increases. The role of cocoon size varies significantly with parasitoid sex, where female parasitoids show a greater rate of increase in size than males. In contrast, though *N. lecontei* are on average larger than *N. pinetum* (Glover et al., 2023), host species does not moderate the relationship between cocoon size and parasitoid size (i.e., there is no significant interaction between cocoon size and host sawfly species). Therefore, *P. neodiprioni* tend to have larger body sizes when they parasitize larger host cocoons, regardless of host identity. This suggests that *P. neodiprioni* across hosts either share preferential host selection methods to parasitize larger individuals (Lin & Ives, 2003), or *P. neodiprioni* across hosts experience similar resource-related size constraints (Silva-Torres et al., 2009).

The interaction of cocoon size and host sex also has a significant impact on parasitoid size. This was expected, as both *N. lecontei* and *N. pinetum* exhibit strong size-related sexual dimorphism, with males being much smaller than females (Glover et al., 2023). Cocoon size and host sex tend to be highly correlated (Glover et al., 2023), and would therefore be expected to have the identified, interconnected effect on parasitoid size.

The significant roles of host cocoon size and cocoon size by host sex interactions are maintained across all three body segments. In the abdomen region, the cocoon size by parasitoid sex interaction is also significant. Yet, in neither the fully body variation nor any individual body segment’s variation was cocoon size found to significantly vary with host species. In every case, host species continues to significantly or marginally significantly associate with parasitoid size variation, even after accounting for resource variation associated with a particular host’s size. This ultimately suggests that the *P. neodiprioni* species has size variation linked closely to host species itself, not just a particular host individual’s size, making a strong case for parasitoid divergence across host environments.

### *P. neodiprioni* Body Size Across Latitudes

*P. neodiprioni* body size displays one final, notable pattern that must be compared to their hosts. Like *Neodiprion* sawflies, *P. neodiprioni* exhibits clinal variation. On average, *P. neodiprioni* increase in size as latitude increases. This positive body size-latitude relation is well-characterized in biological literature, and it occurs across many taxa (Stillwell, 2015). However, for *P. neodiprioni*, this pattern occurs regardless of host species, which does not match sawfly patterns. Among *Neodiprion* hosts, *N. lecontei* body size increases with latitude on average, but *N. pinetum* body size decreases with latitude (Glover et al., 2023). *P. neodiprioni’s* positive body size cline mirrors the *N. lecontei* body size cline, but it is distinct from the negative *N. pinetum* cline. In addition, while the head and abdomen segments of *P. neodiprioni* increase with increases in latitude overall, parasitoids’ heads from *N. pinetum* increase at a greater rate than those from *N. lecontei,* and parasitoids’ abdomens from *N. lecontei* decrease compared to increases in samples from *N. pinetum*.

The lack of negative latitudinal trends of body size in parasitoids from *N. pinetum* has several possible clauses. *N. lecontei* could be the preferred host of *P. neodiprioni.* Or, *N. lecontei* could be the ancestral host, and *P. neodiprioni* may have more recently host shifted onto *N. pinetum*. In either case, one would expect *P. neodiprioni* size patterns to be more closely associated with *N. lecontei* variation than *N. pinetum* variation. This would lead to a latitudinal cline that more strongly mimics the *N. lecontei* cline. Alternatively, given that *N. lecontei* and *N. pinetum* have some degree of gene flow, as exhibited by their hybrid zone (Bendall et al., 2022), the similarity of these hosts may inhibit true ecological divergence of their parasitoids, such that *P. neodiprioni* may show divergence across hosts but not complete mimicry of clinal variation within each host. Finally, the pattern may be a consequence of our sample size, which has a non-continuous geographic cline, as our samples are incidental by-catch from past *Neodiprion* collections. In the future, sampling *P. neodiprioni* from a wider latitudinal range for both host species will be critical to better understanding the implications of latitude by host species patterns.

## Conclusions

Identification of host-associated body size and facial feature size divergence in *P. neodiprioni* has several major implications. First, our research provides the first indication of ecological speciation in *P. neodiprioni*. Environment-associated morphological variation has often played a key role in supporting ecological speciation (Glover, 2023; Nosil, 2012; Ryan et al., 2007; Schluter, 1996), and our research gives the first evidence that ecologically-driven divergence may occur in this parasitoid. In particular, *P. neodiprioni* across distinct *Neodiprion* environments display clear divergence in body size, as well as some divergences in sexual dimorphism. This supports the idea that different *Neodiprion* species may place different selective pressures on their parasitoids. Sufficiently different *Neodiprion* host environments may ultimately facilitate divergence in their parasitoids.

Second, our analysis of parallel, host-associated body divergence across trophic levels in the *Pinus-Neodiprion-P. neodiprioni* system presents the first suggestion of sequential divergence in this, or any, system of externally feeding insects and their parasitoids (Thompson et al., 2022; Stireman et al, 2006). In sequential divergence, environment-driven divergence in a host species facilitates environment-driven divergence in an associated organism (Hood et al., 2015). In this system, pine environment differences are associated with body size as a magic trait in diverging *Neodiprion* host species (Glover et al., 2023), and *P. neodiprioni* parasitoids now appear to exhibit parallel body size differentiation according to their *Neodiprion* hosts. In addition, *P. neodiprioni*’s facial feature divergence occurs only between parasitoids from more ancestrally diverged *Neodiprion* hosts (Herrig et al., 2024), not between parasitoids from sister species in their hybrid zone (Bendall et al., 2022). The degree of divergence among host species may moderate ecological divergence among parasitoids, further suggesting a divergence cascade.

No other studies of externally feeding insects and their parasitoids have identified indications of cascading divergence (Thompson et al., 2022). While additional evidence is required to conclude that sequential divergence occurs in this system (Hood et al., 2015), this is a major step in characterizing the biodiversity-generating power of sequential divergence in parasitoid wasps. Therefore, our results provide novel evidence for sequential divergence and suggest it is worthwhile to continue considering sequential divergence as a possible biodiversity-generating mechanism in plant-external herbivore-parasitoid systems.

Third, from our morphological investigation of body size variation, we can propose future research on potential mechanisms of sequential divergence and isolation. In particular, body size acts as a magic trait in ecological speciation of host *N. lecontei* and *N. pinetum*, and body size diverges in parallel in their parasitoids. As host-associated divergence notably occurs in the abdominal region, it seems reasonable to consider possible mechanical reproductive isolation between parasitoids from these two hosts. If body size divergence acted to isolate parasitoids from different hosts, it would be a magic trait in *P. neodiprioni*, just like in *Neodiprion*, suggesting that magic traits in particular may cascade from one trophic level to another, facilitating host to parasitoid sequential divergence. This seems highly likely, as *P. neodiprioni* from *N. lecontei* and *N. pinetum* display host-associated body size divergence in a small region outside of the hosts’ hybrid zone, but not within the hosts’ hybrid zone. To confirm this pattern of magic traits in hosts driving magic traits in associates, future research must directly assess size-driven reproductive isolation in *P. neodiprioni* through live mating trials.

Finally, our research contributes to broader evaluations of the species diversity of Hymenoptera (Forbes et al., 2018). It is highly likely that parasitoid wasps have a vast reserve of undescribed biodiversity, but it is difficult to accurately estimate the extent of these unknown parasitoid species. However, if we understand the mechanisms that generate parasitoid biodiversity and the frequency with which they occur across different system types, we can better estimate parasitoid diversity. In particular, if sequential divergence is widespread across insect systems, especially among externally feeding insect systems, it would facilitate multiplicative parasitoid divergence and support parasitoid wasps as a reservoir of biodiversity. Therefore, this initial support for sequential divergence in a plant-external herbivore-parasitoid system provides an exciting impetus to further explore cascading host-associated divergence in similar systems.

## Study funding

This research was supported by The Ohio State University’s Fellowship Office, whose Early Offer Eminent Scholar Fellowship supported the graduate student researcher during this project.

## Acknowledgements

We thank Dr. Catherine Linnen and the Linnen Laboratory at The University of Kentucky for their contribution of by-catch *P. neodiprioni* samples for analyses. We thank Carson Kephart for his work to identify parasitoid samples to the species level. We thank Joshua Kiefer and Michael Zwez for their role in photographing samples and collecting data.

## Conflict of Interest

We have no conflict of interest to declare.

## Data Availability

All data and R scripts will be publicly archived with Zenodo upon journal publication of this manuscript.

## Notes

### Competing Interest Statement

The authors have declared no competing interest.

