## Supplemental Figures and Table for "Body size cascades from *Neodiprion* sawfly hosts to *Perilampid* parasitoids: An investigation of sequential divergence"

**Supplemental Materials**

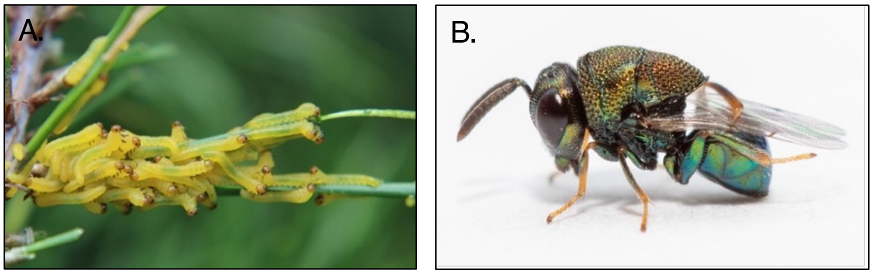


Supplemental Figure 1: Study system. A.) Larval colony of *N. lecontei* feeds on pine host (Bagley, 2016). B.) Adult male *P. neodiprioni* (Ridenbaugh, 2024).
Alt text: Subfigure A shows pine sawfly larvae, and subfigure B shows an adult parasitoid wasp.

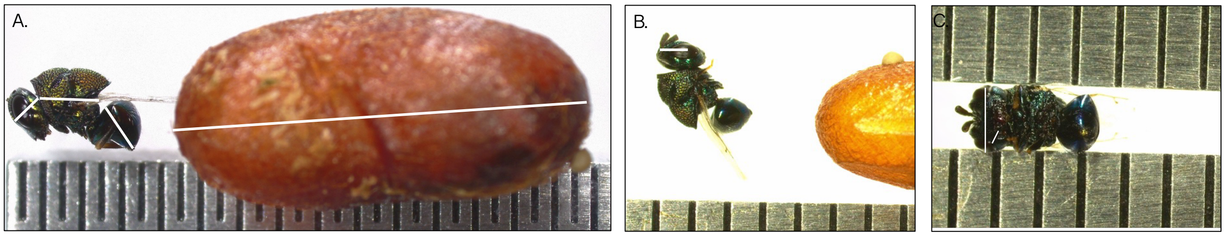


Supplemental Figure 2: A.) Body size measures: Anterior to posterior measurements of parasitoid head, thorax, and abdomen, as well as measurement of *Neodiprion* cocoon. B.) Sexual dimorphism measures: Measurement of eye height. C.) Sexual dimorphism measures: Measurements of two facial features, malar distance and head width.
Alt text: Figures demonstrating examples of body size measurements of a parasitoid wasp. Subfigure A depicts body segment and cocoon measurements, subfigure B uses a white line to demonstrate eye height measure, and subfigure C uses white lines to indicate head width and malar distance.

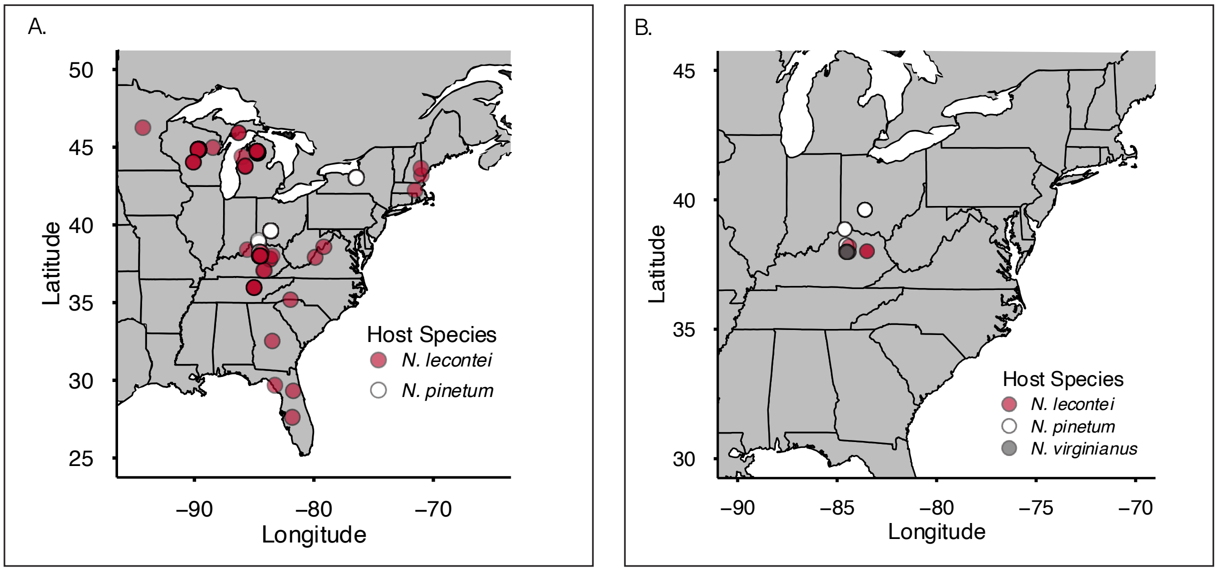


Supplemental Figure 3: A.) *P. neodiprioni* samples in full-range analysis from *N. lecontei* (n=204) and *N. pinetum* (n=103) with each individual sample represented by a single, semi-transparent point. Locations with dense sampling will appear to have a darker color than those with fewer samples. B.) *P. neodiprioni* samples in limited-geography analysis from *N. lecontei* (n=29), *N. pinetum* (n=21), and *N. virginianus* (n=20).
Alt text: Maps of the eastern United States, depicting geographic origins of samples used in study. Subplot A shows samples from the full-range study. Subplot B shows samples from the limited geography study.

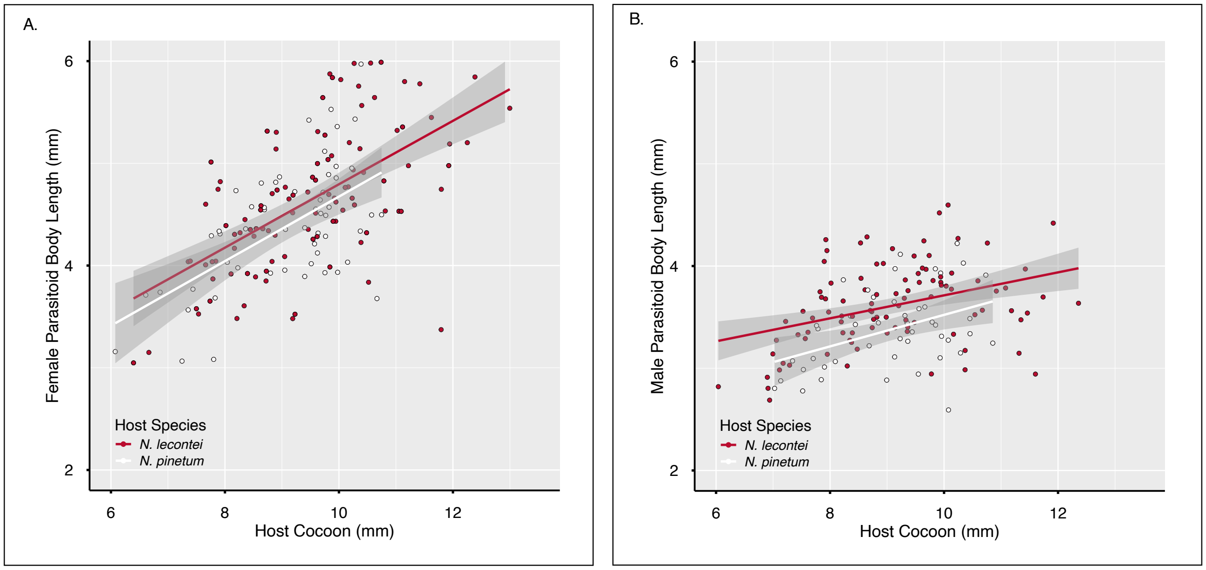


Supplemental Figure 4: A.) Scatterplot with lines of best fit depicting association of sawfly host cocoon size with female parasitoid total body size from *N. lecontei* (red) and *N. pinetum* (white) hosts. B.) Scatterplot with lines of best fit depicting association of sawfly host cocoon size with male parasitoid total body size from *N. lecontei* (red) and *N. pinetum* (white) hosts.
Alt Text: Graphs of scatterplots and lines of best fit, depicting relationship between host cocoon size and parasitoid body size from two sawfly host species. Subplot A depicts female parasitoids, while subplot B depicts male parasitoids.

Supplemental Table 1: Paired comparisons from estimated marginal means analyses of each categorical variable in full-range parasitoid body size analysis (Parasitoid Body Size ~ Host Species + Latitude+ Host Sex + Parasitoid Sex + Cocoon Size + Cocoon Size*Parasitoid Sex + Cocoon Size * Host Sex + smooth(Colony, random effect)). Significant terms are bolded and marginally significant terms are underlined.

| Variable | Comparison | Estimate | Standard Error | df | t-ratio | P-value |
|---|---|---|---|---|---|---|
| **Host Species** | ***N. lecontei - N. pinetum*** | **0.164** | **0.0852** | **246** | **1.925** | **0.0554*** |
| **Parasitoid Sex** | **Female - Male** | **1.02** | **0.052** | **246** | **19.676** | **< 0.001*** |
| **Host Sex** | **Female - Male** | **-0.275** | **0.0927** | **246** | **-2.970** | **0.0033*** |

Supplemental Table 2: Estimated marginal means comparison of body size variation in full-range analysis in relation to the cocoon size by parasitoid sex interaction and cocoon size by host sex interaction. Estimated marginal means results are averaged over levels of the interacting variables, and a 0.95 confidence interval is used. Results are presented as a contrast between levels of the relevant categorical variable. Significant terms are bolded and marginally significant terms are underlined.

| Variable | Comparison | Estimate | Standard Error | df | t-ratio | P-value |
|---|---|---|---|---|---|---|
| **Parasitoid Sex* Cocoon Size** | **Female - Male** | **0.21** | **0.0377** | **246** | **5.562** | **< 0.001*** |
| **Host Sex* Cocoon Size** | **Female - Male** | **-0.205** | **0.0701** | **246** | **-2.929** | **< 0.0037*** |

Supplemental Table 3: Full-range head size multiple regression model ($R^2_{adj}$=0.311, df= 299, p<0.001) and associated type III ANOVA (Parasitoid Head Size ~ Host Species + Latitude+ Host Sex + Parasitoid Sex + Cocoon Size + Latitude*Host Species+ Host Sex*Cocoon Size). Significant terms are bolded and marginally significant terms are underlined.

| Variable | Estimate | Standard Error | t-value | df | Sum Squares | F-value | P-value |
|---|---|---|---|---|---|---|---|
| Intercept | -0.110197 | 0.160680 | -0.686 | 1 | 0.00416 | 0.4703 | 0.49336 |
| **Host Species** | **0.332599** | **0.144423** | **2.303** | **1** | **0.04693** | **5.3036** | **0.02197*** |
| **Latitude** | **0.013254** | **0.003673** | **3.609** | **1** | **0.11525** | **13.0238** | **< 0.001*** |
| **Parasitoid Sex** | **0.039711** | **0.005563** | **7.138** | **1** | **0.45087** | **50.9523** | **< 0.001*** |
| **Cocoon Size** | **0.044231** | **0.007615** | **5.809** | **1** | **0.29858** | **33.7420** | **< 0.001*** |
| Host Sex | 0.128524 | 0.066425 | 1.935 | 1 | 0.03313 | 3.7438 | 0.05395 |
| **Latitude* Host Species** | **-0.008243** | **0.003626** | **-2.273** | **1** | **0.04573** | **5.1674** | **0.02372*** |
| **Cocoon Size*Host Sex** | **-0.016050** | **0.007699** | **-2.085** | **1** | **0.03845** | **4.3454** | **0.03796*** |

Supplemental Table 4: Full-range thorax size generalized additive mixed model ($R^2_{adj}$=0.604, Deviance Explained = 67.5%, n=307) and associated type III ANOVA (Parasitoid Thorax Size ~ Host Species + Latitude+ Host Sex + Parasitoid Sex + Cocoon Size + Cocoon Size*Latitude+ Latitude*Parasitoid Sex + Host Sex*Cocoon Size + Host Sex*Latitude + smooth(Colony, random effect)). Significant terms are bolded and marginally significant terms are underlined.

| Variable | Estimate | Standard Error | t-value | df | Effective df | F-value | P-value |
|---|---|---|---|---|---|---|---|
| **Intercept** | **-8.604299** | **2.309841** | **-3.725** | **NA** | **NA** | **NA** | **0.000241 *** |
| Host Species | 0.044194 | 0.025646 | 1.723 | 1 | NA | 2.970 | 0.086077 |
| **Latitude** | **0.217074** | **0.057747** | **3.759** | **1** | **NA** | **14.130** | **<0.001 *** |
| **Parasitoid Sex** | **0.619499** | **0.225710** | **2.745** | **1** | **NA** | **7.533** | **0.006494*** |
| **Cocoon Size** | **1.095639** | **0.260992** | **4.198** | **1** | **NA** | **17.623** | **< 0.001*** |
| Host Sex | -0.581907 | 0.377432 | -1.542 | 1 | NA | 2.377 | 0.124393 |
| **Cocoon Size*Latitude** | **-0.023047** | **0.006518** | **-3.536** | **1** | **NA** | **12.504** | **< 0.001*** |
| Latitude * Parasitoid Sex | -0.010736 | 0.005560 | -1.931 | 1 | NA | 3.729 | 0.054596 |
| **Cocoon Size*Host Sex** | **-0.051604** | **0.022092** | **-2.336** | **1** | **NA** | **5.456** | **0.020289*** |
| **Latitude*Host Sex** | 0.023507 | 0.008352 | **2.814** | **1** | **NA** | **7.921** | **005274 *** |

| | | | | | | | |
|---|---|---|---|---|---|---|---|
| **Colony** | **NA** | **NA** | **NA** | **NA** | **45.83** | **1.037** | **< 0.001*** |

Supplemental Table 5: Full-range abdomen size generalized additive mixed model ($R^2_{adj}$=0.785, Deviance Explained = 81.9%, n=307) and associated type III ANOVA (Parasitoid Abdomen Size~ Host Species + Latitude+ Host Sex + Parasitoid Sex + Cocoon Size + Cocoon Size*Parasitoid Sex + Latitude* Host Species + smooth(Colony, random effect)). Significant terms are bolded and marginally significant terms are underlined.

| Variable | Estimate | Standard Error | t-value | df | Effective df | F-value | P-value |
|---|---|---|---|---|---|---|---|
| Intercept | -0.894634 | 0.463208 | -1.931 | NA | NA | NA | 0.054533 |
| **Host Species** | **1.035285** | **0.438098** | **2.363** | **1** | **NA** | **5.584** | **0.018865*** |
| **Latitude** | **0.039575** | **0.011081** | **3.571** | **1** | **NA** | **12.756** | **<0.001 *** |
| Parasitoid Sex | -0.125220 | 0.081652 | -1.534 | 1 | NA | 2.352 | 0.126362 |
| **Cocoon Size** | **0.090179** | **0.014809** | **6.089** | **1** | **NA** | **37.081** | **< 0.001*** |
| Host Sex | 0.006846 | 0.019050 | 0.359 | 1 | NA | 7.127 | 0.719603 |
| **Cocoon Size*Parasitoid Sex** | **0.044795** | **0.008760** | **5.113** | **1** | **NA** | **26.147** | **< 0.001*** |
| **Latitude * Host Species** | **-0.025557** | **0.011063** | **-2.310** | **1** | **NA** | **5.337** | **0.021674*** |
| **Colony** | **NA** | **NA** | **NA** | **NA** | **41.61** | **0.71** | **< 0.001*** |

Supplemental Table 6: Statistical outputs of limited-geography eye height ($R^2_{adj}$=0.3803, df=58, p < 0.001), head width, and malar distance models, with associated type III ANOVAs. Significant terms are bolded and marginally significant terms are underlined.

| Model | Variable | Group | Estimate | Standard Error | t-value | P-value | ANOVA F-Value | ANOVA P-value |
|---|---|---|---|---|---|---|---|---|
| Eye Height | **Intercept** | **NA** | **0.891022** | **0.010581** | **84.208** | **< 0.001*** | **7091.0356** | **< 0.001*** |
| | Host Species | Species 1 | -0.002097 | 0.015233 | -0.138 | 0.891 | 0.0220 | 0.97827 |
| | | Species 2 | -0.000903 | 0.015045 | -0.060 | 0.952 | 0.0220 | 0.97827 |
| | **Parasitoid Sex** | **Sex 1** | **0.061000** | **0.010581** | **5.765** | **< 0.001*** | **33.2346** | **< 0.001*** |
| | **Host Species * Parasitoid Sex** | **Species 1 * Sex 1** | **-0.023875** | **0.015233** | **-1.567** | **0.122** | **3.5083** | **0.03645*** |
| | | **Species 2 * Sex 1** | **-0.014547** | **0.015045** | **-0.967** | **0.338** | **3.5083** | **0.03645*** |
| Head Width | **Intercept** | **NA** | **1.6442252** | **0.01358443** | **121.03747** | **0.0000** | **NA** | **< 0.001*** |
| | Host Species | Species 1 | -0.0075036 | 0.01873188 | -0.40058 | 0.6910 | NA | 0.91540 |
| | | Species 2 | 0.0070697 | 0.02309129 | 0.30617 | 0.7612 | NA | 0.91540 |
| | **Parasitoid Sex** | **Sex 1** | **0.1568500** | **0.01355075** | **11.57501** | **0.0000** | **NA** | **< 0.001*** |
| | **Host Species * Parasitoid Sex** | **Species 1 * Sex 1** | **-0.0361990** | **0.01867833** | **-1.93802** | **0.0662** | **NA** | **0.01116*** |
| | | **Species 2 * Sex 1** | **0.0028937** | **0.02303565** | **0.12562** | **0.9012** | **NA** | **0.01116*** |
| Malar Dist. | **Intercept** | **NA** | **0.43049** | **0.01076** | **40.016** | **< 0.001*** | **NA** | **< 0.001*** |
| | Host Species | Species 1 | -0.01078 | 0.01429 | -0.754 | 0.455 | NA | 0.4373 |
| | | Species 2 | 0.01985 | 0.01552 | 1.279 | 0.209 | NA | 0.4373 |
| | **Parasitoid Sex** | **Sex 1** | **0.06632** | **0.01020** | **6.504** | **< 0.001*** | **NA** | **< 0.001*** |

Supplemental Table 7: Statistical outputs of limited-geography total body, head, thorax, and abdomen models, with associated type III ANOVAs. Significant terms are bolded and marginally significant terms are underlined.

| Model | Variable | Group | Estimate | Standard Error | df | t-value | P-value | ANOVA $\chi^2$ | ANOVA P-value |
|---|---|---|---|---|---|---|---|---|---|
| Total Body | **Intercept** | **NA** | **3.68281** | **0.06544** | **26.37178** | **56.281** | **< 0.001*** | **3167.5371** | **< 0.001*** |
| | Host Species | Species 1 | -0.07576 | 0.09219 | 29.63795 | -0.822 | 0.4178 | 3.4040 | 0.1823 |
| | | Species 2 | -0.09158 | 0.09467 | 22.75516 | -0.967 | 0.3435 | 3.4040 | 0.1823 |
| | **Parasitoid Sex** | **Sex 1** | **0.52434** | **0.06486** | **33.80529** | **8.084** | **< 0.001*** | **65.3543** | **< 0.001*** |
| | Host Species * Parasitoid Sex | Species 1 * Sex 1 | -0.16469 | 0.09055 | 46.93613 | -1.819 | 0.0753 | 3.5787 | 0.1671 |
| | | Species 2 * Sex 1 | 0.04789 | 0.09427 | 25.57117 | 0.508 | 0.6158 | 3.5787 | 0.1671 |
| Head | **Intercept** | **NA** | **0.821267** | **0.01194** | **23.177072** | **68.757** | **< 0.001** | **4727.5844** | **< 0.001*** |
| | Host Species | Species 1 | 0.019538 | 0.01685 | 25.862715 | 1.159 | 0.2569 | 1.4629 | 0.48121 |
| | | Species 2 | -0.015599 | 0.01724 | 19.396666 | -0.905 | 0.3767 | 1.4629 | 0.48121 |
| | **Parasitoid Sex** | **Sex 1** | **0.061014** | **0.01188** | **31.436024** | **5.134** | **< 0.001*** | **26.3597** | **< 0.001*** |
| | Host Species * Parasitoid Sex | Species 1 * Sex 1 | -0.033296 | 0.01668 | 45.914537 | -1.996 | 0.0519 | 4.7651 | 0.09232 |
| | | Species 2 * Sex 1 | 0.005053 | 0.01720 | 22.339714 | 0.294 | 0.7716 | 4.7651 | 0.09232 |
| Thorax | **Intercept** | **NA** | **1.40209** | **0.02815** | **36.75680** | **49.815** | **< 0.001*** | **2481.5137** | **< 0.001*** |
| | Host Species | Species 1 | -0.02408 | 0.03880 | 40.17018 | -0.621 | 0.5383 | 0.6372 | 0.72716 |

| | | | | | | | | | |
|---|---|---|---|---|---|---|---|---|---|
| | | Species 2 | -0.00444 | 0.04124 | 35.64893 | -0.108 | 0.9149 | 0.6372 | 0.72716 |
| | **Parasitoid Sex** | **Sex 1** | **0.12458** | **0.02435** | **54.92942** | **5.116** | **< 0.001*** | **26.1721** | **< 0.001*** |
| | **Host Species * Parasitoid Sex** | **Species 1 * Sex 1** | **-0.03566** | **0.03296** | **59.84030** | **-1.082** | **0.2837** | **6.3685** | **0.04141*** |
| | | **Species 2 * Sex 1** | **-0.05623** | **0.03352** | **60.12311** | **-1.677** | **0.0986** | **6.3685** | **0.04141*** |
| Abdomen | **Intercept** | **NA** | **1.47924** | **0.04077** | **17.89238** | **36.287** | **< 2e-16** | **1316.7228** | **< 0.001*** |
| | Host Species | Species 1 | -0.09386 | 0.05415 | 17.93474 | -1.733 | 0.100 | 4.9712 | 0.08327 |
| | | Species 2 | -0.02111 | 0.05854 | 13.75395 | -0.361 | 0.724 | 4.9712 | 0.08327 |
| | **Parasitoid Sex** | **Sex 1** | **0.29968** | **0.03917** | **41.24648** | **7.651** | **1.94e-09** | **58.5370** | **< 0.001*** |

Supplemental Table 8: Estimated marginal means analyses of explanatory variables in total body, head, thorax, and abdomen models in the limited-geography range. For interaction terms of total body, head, and thorax models, pairwise comparisons of the parasitoids' body size traits considered sawfly host species within each parasitoid sex grouping, and all comparisons used a Tukey adjustment. For interpretative clarity, variables that are included in a significant interaction term are only presented in the interaction term's estimated marginal means analysis, not each individual variable's estimated marginal means analysis. The estimated marginal means analysis of abdomen length considered the two explanatory variables seperately, due to the non-significant interaction term. Significant terms are bolded and marginally significant terms are underlined.

| Model | Variable | Group | Contrast | Estimate | Standard Error | df | t-ratio | P-value |
|---|---|---|---|---|---|---|---|---|
| Total Body | Parasitoid Sex* Host Species | Female | *N. lecontei – N. pinetum* | -0.1968 | 0.212 | 6.43 | -0.928 | 0.6430 |
| | | **Female** | ***N. lecontei – N. virginianus*** | **-0.5246** | **0.196** | **26.45** | **-2.680** | **0.0325*** |
| | | Female | *N. pinetum - N. virginianus* | -0.3278 | 0.233 | 9.99 | -1.407 | 0.3739 |
| | | Male | *N. lecontei – N. pinetum* | 0.2284 | 0.281 | 52.02 | 0.814 | 0.6961 |
| | | Male | *N. lecontei – N. virginianus* | 0.0384 | 0.259 | 44.06 | 0.148 | 0.9880 |
| | | Male | *N. pinetum - N. virginianus* | -0.1900 | 0.256 | 41.96 | -0.743 | 0.7397 |
| Head | Parasitoid Sex* Host Species | Female | *N. lecontei – N. pinetum* | -0.00321 | 0.0397 | 4.63 | -0.081 | 0.9964 |
| | | Female | *N. lecontei – N. virginianus* | -0.03806 | 0.0363 | 24.50 | -1.048 | 0.5545 |
| | | Female | *N. pinetum - N. virginianus* | -0.03485 | 0.0431 | 8.05 | -0.809 | 0.7083 |
| | | Male | *N. lecontei – N. pinetum* | 0.07349 | 0.0519 | 52.32 | 1.415 | 0.3407 |

| | | | | | | | | |
|---|---|---|---|---|---|---|---|---|
| | | Male | *N. lecontei – N. virginianus* | 0.08502 | 0.0479 | 44.01 | 1.775 | 0.1897 |
| | | Male | *N. pinetum - N. virginianus* | 0.01153 | 0.0471 | 42.83 | 0.245 | 0.9675 |
| Thorax | Parasitoid Sex* Host Species | Female | *N. lecontei – N. pinetum* | 0.000928 | 0.0783 | 42.8 | 0.012 | 0.9999 |
| | | Female | *N. lecontei – N. virginianus* | -0.180140 | 0.0793 | 37.2 | -2.271 | 0.0725 |
| | | Female | *N. pinetum - N. virginianus* | -0.181068 | 0.0916 | 39.2 | -1.977 | 0.1314 |
| | | Male | *N. lecontei – N. pinetum* | -0.040205 | 0.1010 | 56.8 | -0.397 | 0.9169 |
| | | Male | *N. lecontei – N. virginianus* | 0.074947 | 0.1020 | 47.0 | 0.737 | 0.7428 |
| | | Male | *N. pinetum - N. virginianus* | 0.115153 | 0.0982 | 42.2 | 1.172 | 0.4760 |
| Abdomen | **Parasitoid Sex** | **NA** | **Female - Male** | **0.599** | **0.0814** | **50.1** | **7.366** | **< 0.001*** |
| | Host Species | NA | *N. lecontei – N. pinetum* | -0.0727 | 0.1040 | 24.0 | -0.697 | 0.7678 |
| | Host Species | NA | *N. lecontei – N. virginianus* | -0.2088 | 0.0979 | 31.7 | -2.134 | 0.0990 |
| | Host Species | NA | *N. pinetum - N. virginianus* | -0.1361 | 0.1070 | 25.3 | -1.270 | 0.4249 |
